# Regulated alternative splicing of Dscam2 is required for somatosensory circuit wiring

**DOI:** 10.1101/2023.03.01.530539

**Authors:** Samantha E. Galindo, Grace Ji-eun Shin, S. Sean Millard, Wesley B. Grueber

**Affiliations:** Department of Genetics and Development, Vagelos College of Physicians and Surgeons, Columbia University, New York, NY 10032, USA; Mortimer B. Zuckerman Mind Brain Behavior Institute, Columbia University, New York, NY 10027, USA; School of Biomedical Sciences, The University of Queensland, Brisbane, Australia; Department of Physiology and Cellular Biophysics, Vagelos College of Physicians and Surgeons, Columbia University, New York, NY 10032, USA; Department of Neuroscience, Vagelos College of Physicians and Surgeons, Columbia University, New York, NY 10027, USA

**Keywords:** Down syndrome cell adhesion molecule, Dscam2, axon targeting, *Drosophila*, neural circuit, somatosensory neurons, nociception

## Abstract

Axon and dendrite placement and connectivity is guided by a wide range of secreted and surface molecules in the developing nervous system. Nevertheless, the extraordinary complexity of connections in the brain requires that this repertoire be further diversified to precisely and uniquely regulate cell-cell interactions. One important mechanism for molecular diversification is alternative splicing. *Drosophila Down syndrome cell adhesion molecule (Dscam2*) undergoes cell type-specific alternative splicing to produce two isoform-specific homophilic binding proteins. Regulated alternative splicing of *Dscam2* is important for dendrite and axon patterning, but how this translates to circuit wiring and animal behavior is not well understood. Here, we examined the role of cell-type specific expression of *Dscam2* isoforms in regulating synaptic partner selection in the larval somatosensory system. We found that synaptic partners in the nociceptive circuit express different Dscam2 isoforms. Forcing synaptic partners to express a common isoform resulted in nociceptive axon patterning defects and attenuated nocifensive behaviors, indicating that a role for Dscam2 alternative splicing is to ensure that synaptic partners do not express matching isoforms. These results point to a model in which regulated alternative splicing of *Dscam2* across populations of neurons restricts connectivity to specific partners and prevents inappropriate synaptic connections.

## Introduction

Nervous system function is determined by the organization of a tremendous number of neurons into precisely wired neural circuits. During circuit wiring, neurons recognize and respond to molecular cues presented on growth substrates and neuronal membranes. The outcome of recognition events can vary widely, from repulsion to growth and adhesion (Tessier-Lavigne and Goodman, 1996). Ultimately, the outcome of these many events is to register presynaptic axons and postsynaptic dendrites at a shared meeting place, an important early step in connection specificity. Within this spatially restricted area, highly specific synaptic connectivity can be achieved through mutual recognition mediated by cell recognition molecules (CRMs) (Yogev and Shen, 2014).

CRMs can promote synaptic specificity by providing molecular recognition codes that instruct connectivity. Given the vast requirement for recognition selectivity in the construction of the nervous system, mechanisms must exist to diversify the repertoire of neural recognition molecules. Diversification can be achieved through alternative splicing, as has been shown in *Drosophila* for *Down syndrome cell adhesion molecule (Dscam*) genes. Extreme diversification at the *Dscam1* locus can generate over 38,000 distinct isoforms (Schmucker et al., 2000). Stochastic use of Dscam1 isoforms produces recognition codes for individual neurons and promotes self-avoidance (Hughes et al., 2007; Matthews et al., 2007), but is ill-suited for deterministic synaptic specificity. By contrast, a more restricted and tightly regulated alternative splicing, as is seen at the Dscam2 locus may be well suited for roles in connection specificity (Kerwin et al., 2018; Lah et al., 2014; Li and Millard, 2019). Given the extensive diversification of recognition molecules, it is conceivable that connection specificity would be determined by a dedicated code that is specific to each connected pair. However, strategies for connectivity that work for one cell pair may also be reused in a circuit such that populations of neurons that share similar connectivity share CRM recognition codes. Such recognition codes could promote pairing between appropriate partners, or, conversely, inhibit synapse formation between inappropriate partners.

*Down syndrome cell adhesion molecule 2 (Dscam2*) undergoes cell type-specific alternative splicing to encode two isoforms, Dscam2A and Dscam2B. Dscam2A and 2B differ by a single extracellular immunoglobulin domain and bind in an isoform-specific homophilic manner (Millard et al., 2007). Dscam2 controls tiling, dendrite and axon patterning, and synaptic specificity in the visual system (Kerwin et al., 2018; Lah et al., 2014; Millard et al., 2007; Millard et al., 2010; Tadros et al., 2016) and regulates synaptic strength in motor neurons in an isoform-specific manner (Odierna et al., 2020). Given the repulsive outcome of Dscam2-mediated cell-cell interactions, Dscam2 may promote cell type-specific avoidance between non-partners, complementing mechanisms of instructive synaptic matching. While the role of Dscam2 has been studied extensively at the level of cell-cell contact and synapse formation, it is not known whether regulated Dscam2 isoform expression contributes to the wiring of functional neural circuitries.

Here, we investigated the role of cell type-specific Dscam2 isoform expression in mediating interactions between pre- and postsynaptic partners in the *Drosophila* larval somatosensory system. The larval body wall is innervated by dendritic arborization (da) sensory neurons, which are categorized into four classes (class I-IV, or cI-cIV) based on morphology and function (**Figure 1A**) (Grueber et al., 2002; Grueber et al., 2007). cIV nociceptive neurons detect noxious mechanical and thermal stimuli (Hwang et al., 2007a) and are the most comprehensively studied class. Because of powerful behavioral assays (Hwang et al., 2007a), tools for specific labeling and manipulation, and the recent identification of all second-order neurons and many higher-order circuit components (Burgos et al., 2018; Gerhard et al., 2017; Hu et al., 2017; Kaneko et al., 2017; Ohyama et al., 2015; Takagi et al., 2017; Yoshino et al., 2017), the larval nociceptive system is an excellent model for studies of the development and function of neural circuits. We find that distinct Dscam2 isoforms are expressed in nociceptive sensory neurons and downstream interneurons. We demonstrate that cell type-specific alternative splicing of Dscam2 regulates interactions between synaptic partners in this circuit and is critical for nociceptive sensory axon patterning and function. Our data support the conclusion that non-matching isoform expression in pre- and postsynaptic partners is important for sensory to interneuron synaptic connectivity and function of nociceptive circuitry in *Drosophila*. These findings provide an example of how regulated alternative splicing contributes to the expansion of recognition codes during nociceptive circuit formation.

**Figure 1.**
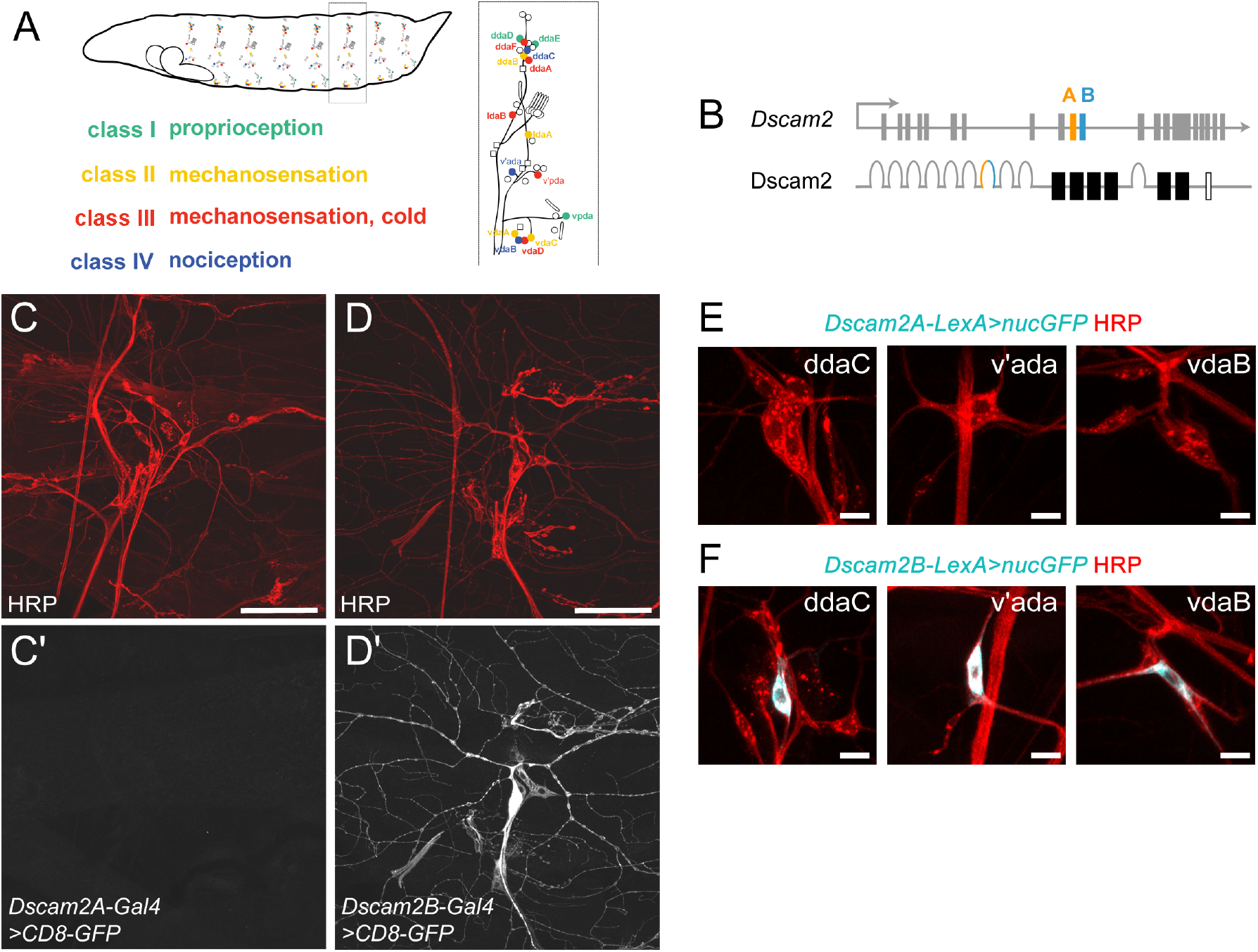
Dscam2B is expressed in class IV nociceptive sensory neurons. (A) Diagram of larval dendritic arborization neurons. (B) Dscam2 gene (top) and protein (bottom). Constant exons are drawn as gray rectangles, variable exon 10 is drawn as orange (10A) or blue (10B) rectangle. Gray horseshoes represent constant Immunoglobulin (Ig) domains, colored horseshoe indicates variable Ig domain, solid black rectangles represent Fibronectin domains, black rectangle (no fill) represents transmembrane domain. (C-D’) Maximum intensity projection of dorsal cluster sensory neurons labeled by anti-HRP antibody (C, D) or anti-GFP antibody (C’, D’). *Dscam2A-Gal4* (C’) and *Dscam2B-Gal4* (D’) driving *UAS-CD8-GFP* (with anti-GFP stain) show that *Dscam2A-Gal4* is not expressed in sensory neurons and *Dscam2B-Gal4* is expressed in ddaC class IV neurons. (E) *Dscam2A-LexA* expression (cyan) is not observed in class IV cell bodies (labeled by HRP). (F) class IV cell bodies labeled by *Dscam2B-LexA* (cyan) and HRP (red). Scale bars: 50 μm (C-D’) and 10 μm (E-F).

## Results

### Synaptic partners in the larval nociceptive circuit express complementary Dscam2 isoforms

The *Dscam2* locus generates two alternatively spliced isoforms, Dscam2A and Dscam2B (Lah et al., 2014; Millard et al., 2007) (**Figure 1B**). To characterize Dscam2 isoform expression in the larval da sensory neurons, we used isoform-specific Gal4 and LexA splicing reporters (Lah et al., 2014; Tadros et al., 2016) to drive expression of a membrane-targeted or nuclear envelope-targeted GFP and characterize endogenous Dscam2 isoform expression in third instar larvae. In flies bearing these constructs, Gal4 or LexA are knocked into the variable region of *Dscam2* so that the expression pattern of these proteins reflects the endogenous expression patterns and levels of *Dscam2* isoforms. We did not detect Dscam2A-Gal4 or Dscam2A-LexA expression in body wall neurons whereas Dscam2B-Gal4 and Dscam2B-LexA were strongly expressed in class IV da neurons (**Figure 1C-F**).

We also examined Dscam2 isoform expression in the larval CNS in somatosensory interneurons. Consistent with published results (Odierna et al., 2020), isoform reporters showed broad expression in the brain and ventral nerve cord (**Figure 2, A-D**). To investigate whether isoform choice is regulated in nociceptive circuits, we focused on interneurons that are direct postsynaptic partners of cIV neurons, termed Down-and-Back (DnB) neurons and Basin neurons (**Figure 2E, F**) (Burgos et al., 2018; Ohyama et al., 2015). DnB neurons receive the majority of their synaptic input from cIV neurons (Burgos et al., 2018). Basin neurons comprise four sub-populations of neurons (Basin1-4), which receive either significant input (Basin 2,4) or minor input (Basin 1,3) from cIV neurons (Ohyama et al., 2015). We used Dscam2A- and Dscam2B-LexA reporters to drive expression of *LexAop-CD8-GFP*, and simultaneously visualized DnB neurons (labeled by *412-Gal4*) or Basin neurons (labeled by *72F11-Gal4*) with *UAS-CD8-mcherry* (**Figure 2G**). We found that almost all (98%, n=62/63) Basin neurons we examined and most DnB neurons (74%, n=35/47) expressed *Dscam2A-LexA*. In contrast, we did not observe *Dscam2B-LexA* expression in either Basin or DnB neurons (n=0/78, n=0/47, respectively) (**Figure 2H**). These data show that Dscam2 isoforms are expressed in a tissue-specific and cell-type-specific manner in the larval nociceptive circuit, with first-order synaptic partners expressing different isoforms than cIV neurons (**Figure 2I**).

**Figure 2.**
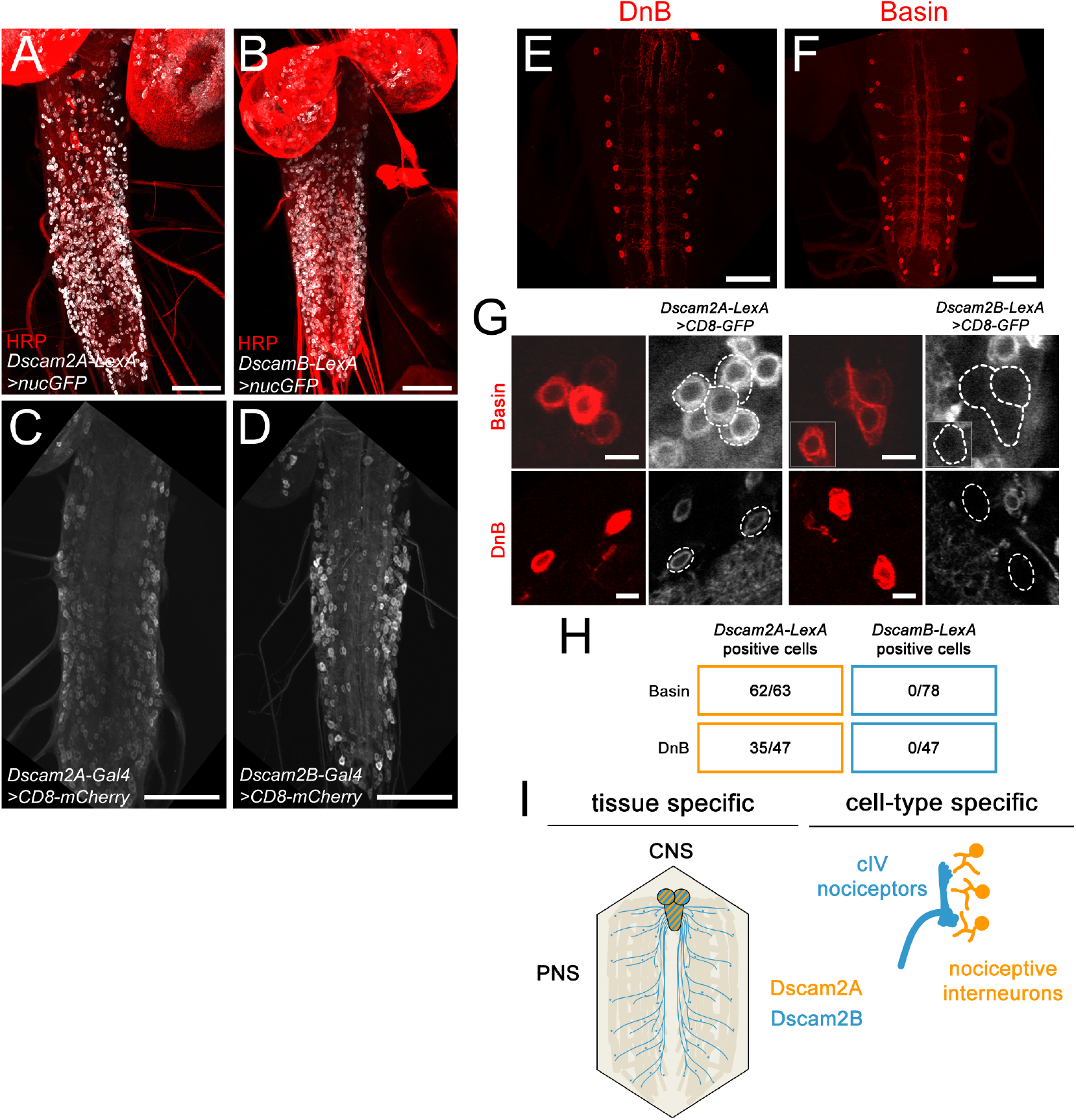
Dscam2A is expressed in nociceptive interneurons. (A-B) Maximum intensity projections (Max IPs) of *Dscam2A-LexA* (A) and *Dscam2B-LexA* (B) expression (gray) in the CNS, co-labeled with anti-HRP antibody (red). (C-D) Max IPs of *Dscam2A-Gal4* (C) and *Dscam2B-Gal4* (D) expression in the VNC. (E-F) Max IPs of DnB neurons (E) and Basin neurons (F) labeled by *412-Gal4* and *72F11-Gal4*, respectively. (G) Single-plane images of *Dscam2A-LexA* and *Dscam2B-LexA*-driven CD8-GFP. *Dscam2A-LexA*, but not *Dscam2B-LexA* is expressed in DnB and Basin^1-4^ neurons (labeled by CD8-mCherry). (H) Number of cells examined per cell type and isoform. Three larvae were examined for each cell type/isoform. (I) Summary of Dscam2 isoform expression in larval nervous system. Scale bars: (A-F) 50 μm (G) 5 μm

To explore the basis for isoform-specific Dscam2 expression in class IV neurons, we examined whether misexpression of Knot, a COE transcription factor that is expressed in cIV neurons and controls their identity (Crozatier and Vincent, 2008; Hattori et al., 2007; Jinushi-Nakao et al., 2007), can induce Dscam2B expression in other da neuron classes. We expressed Knot under the control of pan-multidendritic (md) sensory neuron driver *109(2)80-Gal4* and used *Dscam2B-LexA* driven myr-GFP as a readout for Dscam2B expression. In control larvae we consistently observed *Dscam2B-LexA* expression in class IV neurons (85% of ddaC cells, 93% of v’ada cells, 40% of vdaB cells) (**Figure 3A, C**). When we forced Knot expression we observed a significant reduction in the number of cIV neurons that expressed *Dscam2B-LexA*, indicating that levels of the Knot transcription factor may modulate transcription of *Dscam2B* in somatosensory neurons (**Figure 3C**). Though we did not observe widespread ectopic expression of *Dscam2B-LexA* in other sensory neurons, we observed a significant increase in *Dscam2B-LexA* expression in td sensory neurons that innervate the tracheal system (43% of neurons) (**Figure 3B, C**). The basis for this selective Knot-induced expression is not known, however, our result suggests that Knot is sufficient to induce ectopic *Dscam2B-LexA* expression in selected sensory neurons.

**Figure 3.**
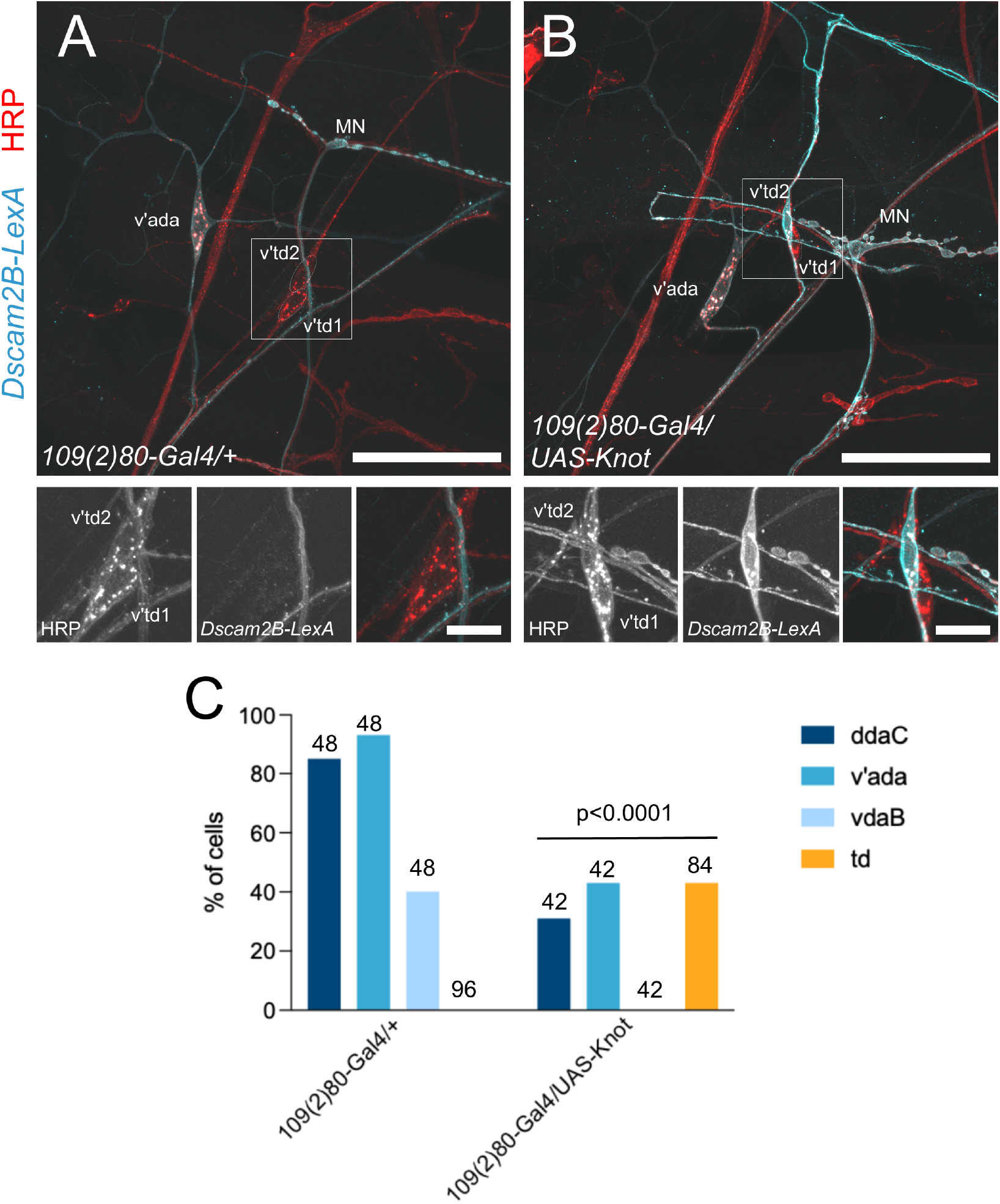
A cIV-specific transcription factor can ectopically activate Dscam2B-LexA in sensory neurons. (A-B) Max IP of Dscam2B-LexA-driven myrGFP (cyan), co-labeled with HRP (red) in lateral body wall of control (A) and larvae in which the Knot transcription factor is mis-expressed in md neurons (B). Tracheal dendrite (td) neurons (v’td1) express Dscam2B-LexA when they ectopically express Knot, but not in control. (C) Quantification of cells examined that express Dscam2B-LexA in A and B. Number of cells (n) examined are shown above bars. p<0.0001 for control vs. Knot ME (ddaC, v’ada, vdaB, td) as determined by two-tailed Fisher’s exact test. 8 control larvae and 8 Knot ME were examined. Scale bars: 50 μm in A and B, 10μm in cell body insets.

### Restricting Dscam2 isoform diversity between synaptic partners in the nociceptive circuit disrupts wiring

We next asked whether Dscam2 isoform diversity is important for nociceptive circuit wiring by examining nociceptive axon patterning in larvae that lack alternative splicing of Dscam2. We examined two mutants harboring modifications of the endogenous *Dscam2* locus that lack alternative splicing of Dscam2 (Lah et al., 2014). In *Dscam2A* single isoform mutants, the variable region containing exon 10A and 10B is replaced with 10A, such that all Dscam2-expressing cells express Dscam2A under otherwise normal regulatory elements. Conversely, in *Dscam2B* single isoform mutants, the variable region is replaced with exon 10B, forcing all Dscam2 expressing cells to produce only the Dscam2B transcript (**Figure 4A**). We introduced a marker for cIV neurons, *ppk^EGFP^*(Grueber et al., 2003), into homozygous *Dscam2A* and *Dscam2B* backgrounds and examined the axon projections of nociceptive cIV neurons. We found that Dscam2 isoform diversity is not required for guidance of cIV axons to the somatosensory neuropil. However, we observed thin or missing longitudinal axon projections, abnormal accumulation of commissural branches, and some axons that were either mistargeted or missing entirely from the neuropil (**Figure 4B, C**). These defects suggest that alternative splicing of Dscam2 is important for local patterning of cIV axon terminals.

**Figure 4.**
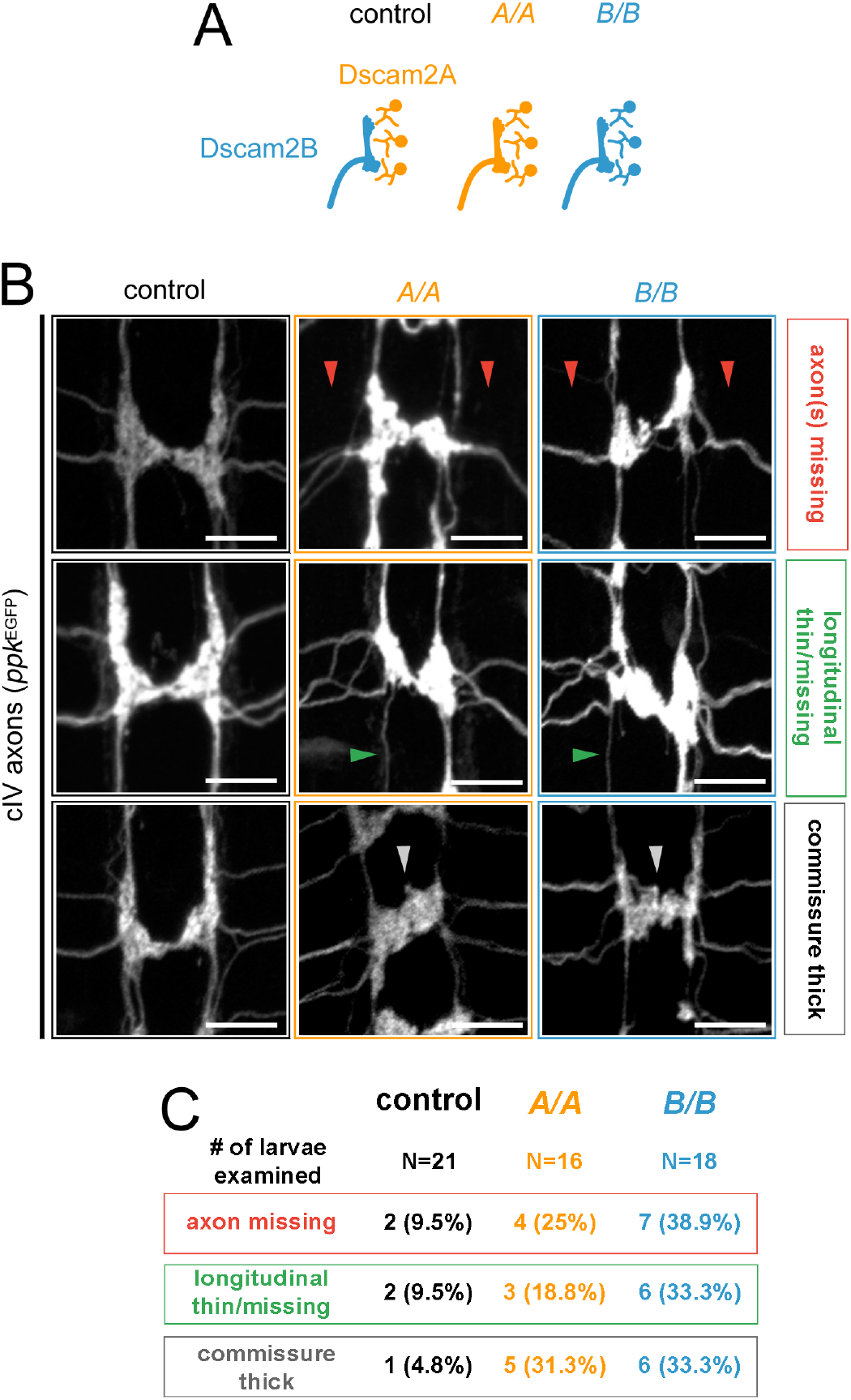
Alternative splicing of Dscam2 is required for nociceptive axon targeting. (A) Schematic of Dscam2 isoform expression in nociceptive sensory and target interneurons in control larvae and Dscam2 single isoform mutants (A/A and B/B). (B) Max IPs of cIV axon terminals (labeled by *ppk^EGFP^*) in control (*w^1118^*) and *Dscam2* single isoform mutant larvae showing patterning defects. Red arrowheads indicate missing axons, green arrowheads indicate missing or thin longitudinal tracts, gray arrowheads indicate thickened commissural axons crossing the midline. (C) Quantification of number of larvae with phenotypes shown in B. Number of larvae (N) examined = 21 (control), 16 (Dscam2A/A), 18 (Dscam2B/B). Scale bars: 10 μm.

In single isoform mutant larvae, the population of Dscam2-expressing cells is composed of a mosaic of cells experiencing isoform-specific loss- and gain-of-function compared to wild-type larvae: cells that express Dscam2B in wild-type animals lack Dscam2B in *Dscam2A* mutant animals and gain expression of Dscam2A, and *vice versa*. Because we observed abnormal axon organization in both *Dscam2A* and *Dscam2B* single isoform homozygotes, a cell-autonomous requirement for Dscam2B in cIV neurons is unlikely to be the sole underlying cause of disrupted axon patterning. Alternatively, cIV axon targeting defects in single isoform mutants may arise because all Dscam2-expressing cells are expressing a common isoform, which would support at least partial non-cell autonomous effects of Dscam2 on cIV neurons. To test this hypothesis, we examined the role of isoform diversity between cIV sensory neurons and their synaptic targets, DnB neurons and Basin neurons. We expressed *UAS-Dscam2B* in DnB and Basin interneurons under the control of *412-Gal4* and *72F11-Gal4*, and visualized cIV neurons using *ppk-CD4-tdTomato* (**Figure 5A**). We found that forcing nociceptive synaptic partners to express a common isoform resulted in cIV targeting defects. Specifically, cIV axons that shared a common isoform with their targets lacked longitudinal axons (**Figure 5B, C**). This defect was specific to shared isoform expression between pre- and postsynaptic partners, as over-expressing Dscam2A in DnB and Basin neurons (their endogenous isoform) did not impact cIV axon targeting. These results suggest that isoform diversity between cIV neurons and their targets is required for proper cIV axon targeting.

**Figure 5.**
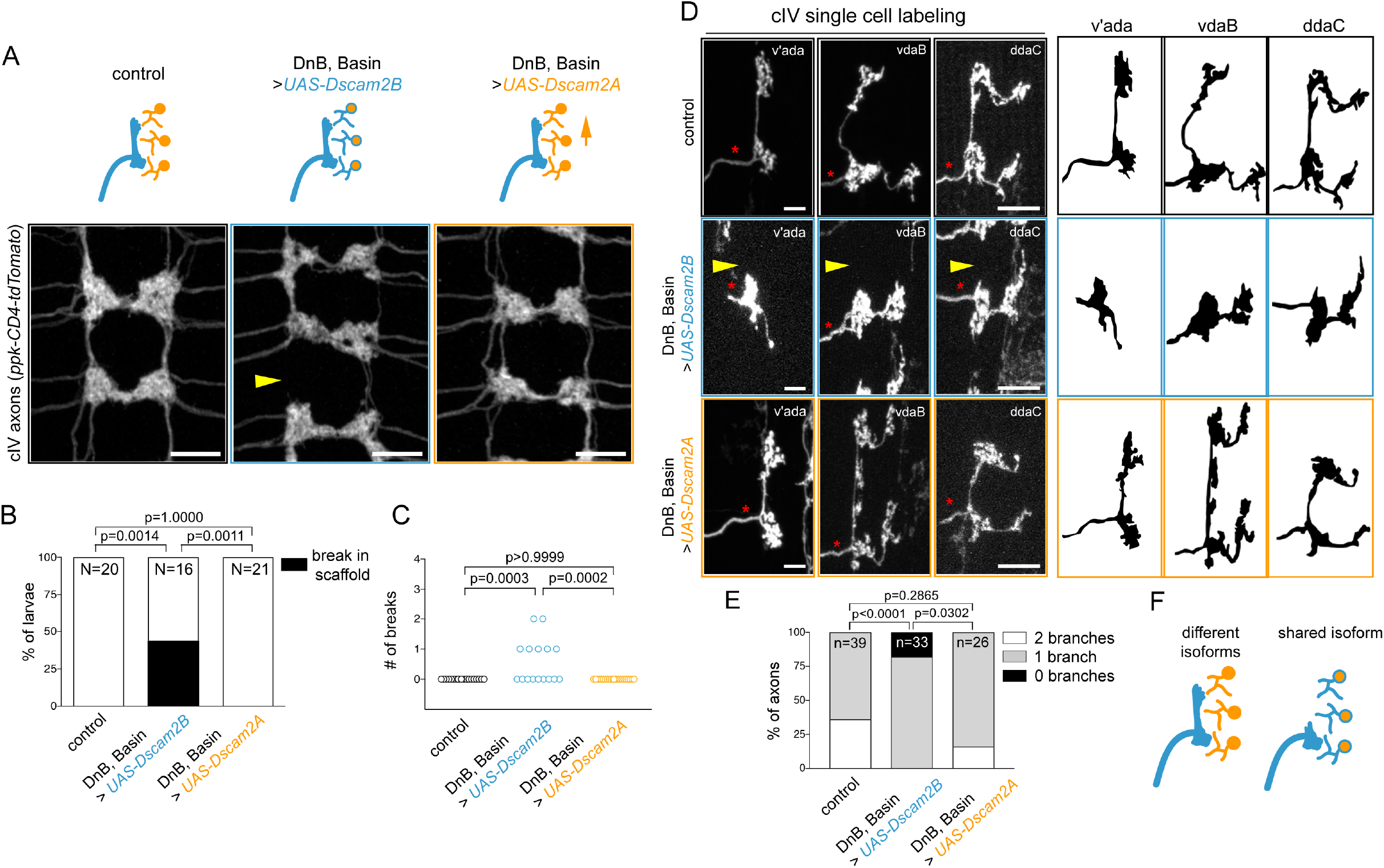
Restricting Dscam2 isoform diversity between synaptic partners in the nociceptive circuit disrupts axon patterning. (A) Top: Schematic of Dscam2 isoform mis- and over-expression paradigms. Orange interneurons express Dscam2A (endogenous), blue sensory neurons express Dscam2B (endogenous), orange interneurons with blue outline express Dscam2A (endogenous) and misexpress Dscam2B, orange interneurons with orange arrow express Dscam2A (endogenous) and over-express Dscam2A. Bottom: Max IP images of cIV axon terminals from adjacent neuromeres in control larvae, larvae in which cIV, DnB and Basin neurons share a common isoform (Dscam2B), and larvae in which DnB and Basin interneurons overexpress Dscam2A to control for potential effects of isoform overexpression. Note that longitudinal axons are missing when DnB and Basin neurons express Dscam2B (yellow arrowhead) but not Dscam2A. (B-C) Quantification of cIV axon targeting defects shown in A. B shows % of larvae that had at least one break in the cIV axon scaffold. Control 0% (0/20); Dscam2B 43.75% (7/16); Dscam2A 0% (0/21) where N indicates number of larvae. Control vs. Dscam2B, p=0.0014; Control vs. Dscam2A, p=1.0000); Dscam2A vs. Dscam2B, p=0.0011, as determined by two-tailed Fisher’s exact test with Bonferroni correction for multiple comparisons. C shows number of breaks observed in each scaffold. Each dot represents one larva. Control vs. Dscam2B, p=0.0003; Control vs. Dscam2A, p>0.9999; Dscam2A vs. Dscam2B p=0.0002, as determined by Kruskal-Wallis test with Dunn’s multiple comparison test. (D) Max IP images of cIV axon terminals under Dscam2 mis/over-expression strategies outlined in (A). cIV axon single cell labeling was produced using the FLPout technique to achieve sparse labeling. Asterisks indicate branchpoints where the axon bifurcates. Note that longitudinal axons are missing when only when DnB and Basin neurons express Dscam2B (arrowheads), matching endogenous Dscam2B expression in cIV neurons. (E) Quantification of axon terminal defects in E. (n) refers to number of single cIV axons. Kruskal-Wallis with Dunn’s multiple comparisons test was used to compare number of branches per axon between control, Dscam2B misexpression, and Dscam2A overexpression groups. Control vs. DscamB, p<0.0001, control vs. Dscam2A, p=0.2865, Dscam2B vs. Dscam2A, p=0.0302. (F) Schematic of a cIV axon phenotype when DnB and Basin neurons mis-express Dscam2B (colabeled with blue and orange), matching endogenous Dscam2B expression in cIV neurons (blue). Scale bars: (B) 10 μm (E) 5 μm

To understand the basis of cIV axon scaffold defects when DnB and Basin neurons misexpress Dscam2B, we used MultiColor FLPout (Nern et al., 2015) to examine individual cIV axon terminals. When target interneurons were forced to express Dscam2B, 18% of nociceptive axon terminals lacked their typical longitudinal process, compared to 0% for control axon terminals (**Figure 5D-F**). We confirmed that this defect was specific to common isoform expression between cIV neurons and their partners, as overexpressing Dscam2A in DnB and Basin neurons did not result in missing cIV axon branches. These results indicate that perturbing the isoform differences between cIV sensory neurons and their synaptic targets reduces sensory axon branching.

### Disruption of Dscam2 diversity is accompanied by reduction of presynapses

We next investigated whether the disruption of cIV axon branching we observed when isoform diversity is perturbed is accompanied by reduction in presynapses. We examined the distribution of the active zone marker brp-short^cherry^ (Berger-Muller et al., 2013) when pre- and postsynaptic partners expressed endogenous isoforms and when they shared a common isoform via misexpression of *UAS-Dscam2B* in DnB and Basin neurons. We visualized cIV axons and presynaptic sites using *38A10-LexA* (Pfeiffer et al., 2010) to drive a membrane-targeted GFP and Brp-short^cherry^. In control larvae, we observed many Brp-short puncta throughout cIV axon terminals, including a few puncta in the longitudinal axons (**Figure 6A**). In larvae in which cIV, DnB and Basin neurons shared a common isoform, the absence of longitudinal cIV axons was accompanied by a concomitant loss of presynaptic sites in those axons (**Figure 6A**, arrowheads). Furthermore, brp-short signal (as measured by fluorescence) in cIV axons was significantly reduced in larvae in which cIV, DnB, and Basin neurons shared a common isoform compared to controls (**Figure 6B**). These results indicate that perturbing isoform diversity between cIV neurons and their targets results in a reduction in cIV presynaptic sites.

**Figure 6.**
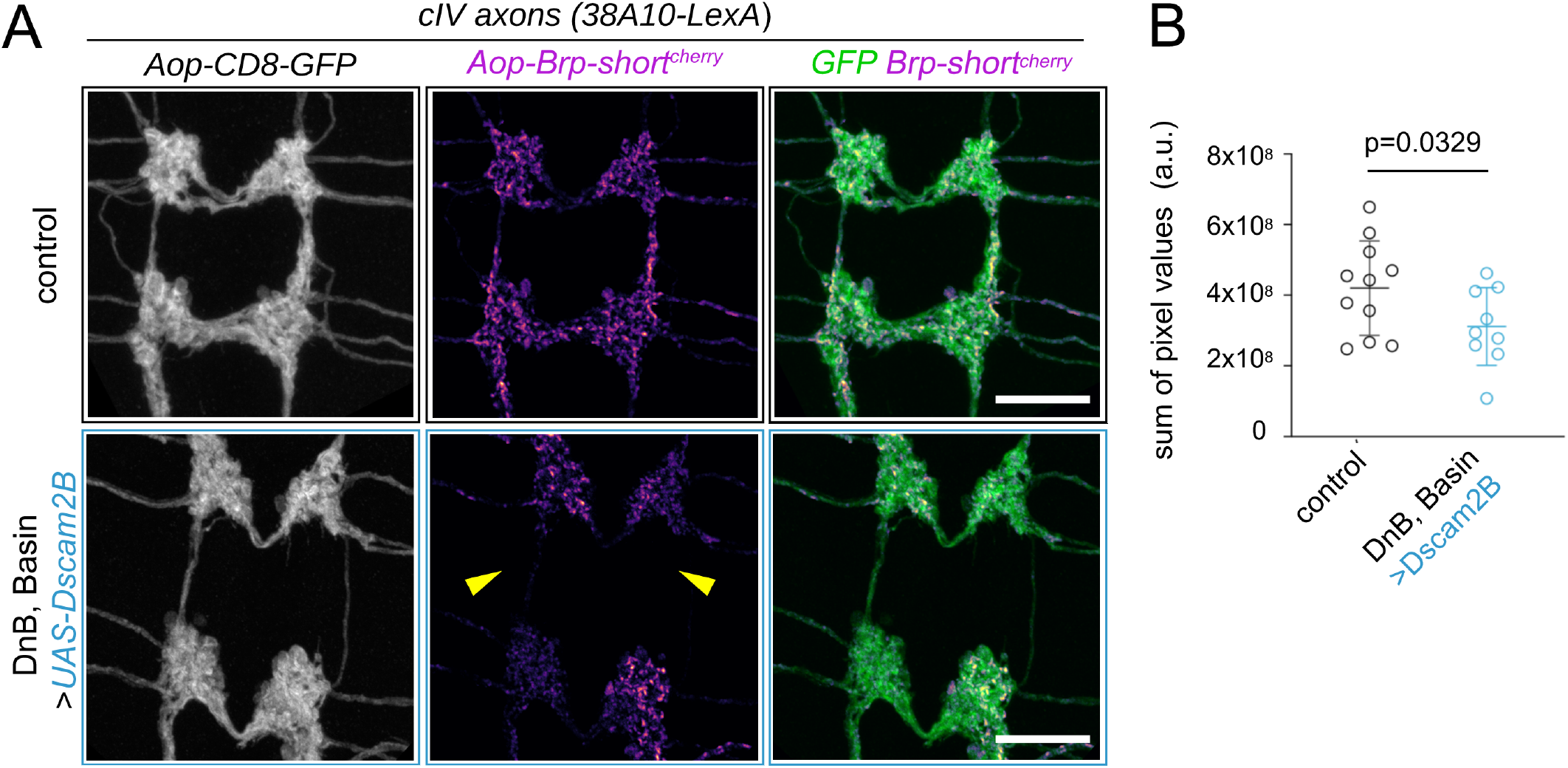
Restricting Dscam2 isoform diversity between synaptic partners leads to a reduction of presynaptic marker. (A) Max IP images of cIV axon terminals and presynaptic sites labeled by CD8-GFP and brp-short^cherry^. Top row shows control cIV axons and bottom row shows cIV axons when DnB and Basin neurons mis-express Dscam2B. (B) Fluorescence (sum of pixel values) in cIV terminals. Plot shows sum of pixel values measured in segments A4-A7. Each dot represents one larva. Error bars indicate ± s.d. Control vs. Dscam2B, p = 0.033, as determined by one-tailed Unpaired t test. Scale bars: 10 μm

### Excess or ectopic Dscam2 causes defasciculation of nociceptive axon terminals

We next examined whether excess Dscam2 in cIV neurons promotes adhesion or repulsion between adjacent axon terminals. Normally, cIV axon terminals form cohesive bundles with branches that either project along the anterior-posterior axis (longitudinal branches) or across the midline (commissural branches) (Grueber et al., 2007). We expressed *UAS-Dscam2A* or *UAS-Dscam2B* in cIV neurons using *ppk^1.9^-Gal4* (Ainsley et al., 2003) in wild-type animals, and visualized cIV axons using *ppk-CD4-tdGFP* (Han et al., 2011). Forced expression of either isoform resulted in prominent defects in axon scaffold organization, most notably axon defasciculation along longitudinal tracts (**Figure 7A-B**). Thus, it appears that excessive amounts of Dscam2 in nociceptive neurons prevent the association between adjacent axon terminal branches, and consequently cause excess spacing between branches.

**Figure 7.**
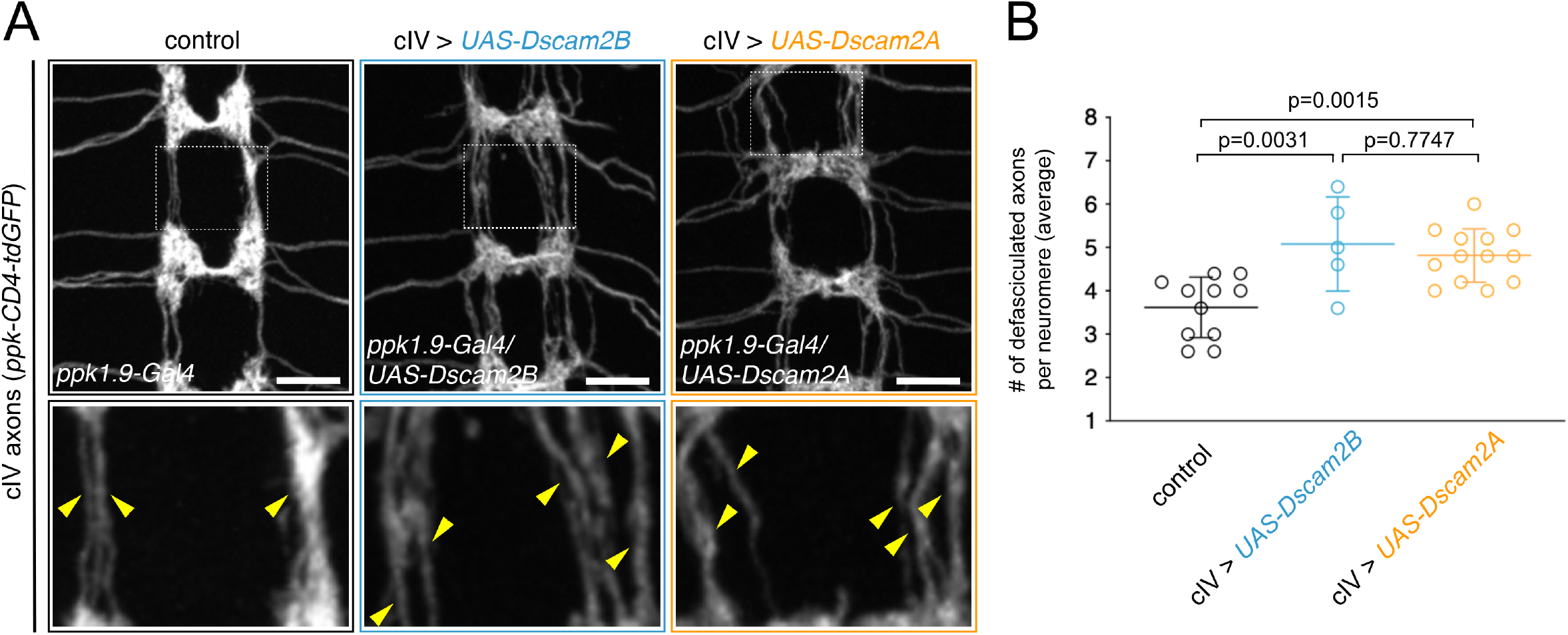
Excess Dscam2 induces segregation between nociceptive axon terminals. (A) Max IP images of control cIV axon terminals (labeled by *ppk-CD4-tdGFP*) compared to cIV axon terminals over-expressing Dscam2B or mis-expressing Dscam2A. Note that longitudinal axon terminals segregate when expressing excess either of Dscam2 isoforms (arrowheads). (B) Quantification of axon defasciculation observed in (A). Neuromeres were scored for total number of longitudinal connectives. Each dot represents the average (A2-A8) for a single larva. Bars show population means ±s.d. control vs. Dscam2B, p=0.0031; control vs. Dscam2A, p=0.0015; Dscam2B vs. Dscam2A, p=0.7747, as determined by ANOVA with Sidak multiple comparison test. Scale bars: 10 μm

### Dscam2 isoform diversity is required for normal nociceptive behavior

We next asked whether regulated alternative splicing of Dscam2 is important for nociceptive circuit function. When presented with noxious thermal stimuli, larvae perform an escape sequence consisting of rapid “corkscrew-like” rolling, followed by fast crawling (Hwang et al., 2007b) (**Figure 8A**). We tested the ability of Dscam2 single isoform mutants to respond appropriately to noxious heat by placing them for 30 seconds on an agar plate heated to a noxious temperature (41 °C) and analyzing rolling behavior. Compared to control (*w^1118^*) larvae, *Dscam2A* single isoform homozygote larvae performed significantly fewer rolls (**Figure 8B**). *Dscam2B* single isoform homozygote larvae did not show any significant difference in number of rolls compared to wild-type, however, the duration of the rolling bout was significantly increased in *Dscam2B* single isoform homozygotes (**Figure 8C**), suggesting that rolling is slowed. Taken together, these results show that regulated alternative splicing of both Dscam2 isoforms is required for efficient nociceptive rolling behavior.

**Figure 8.**
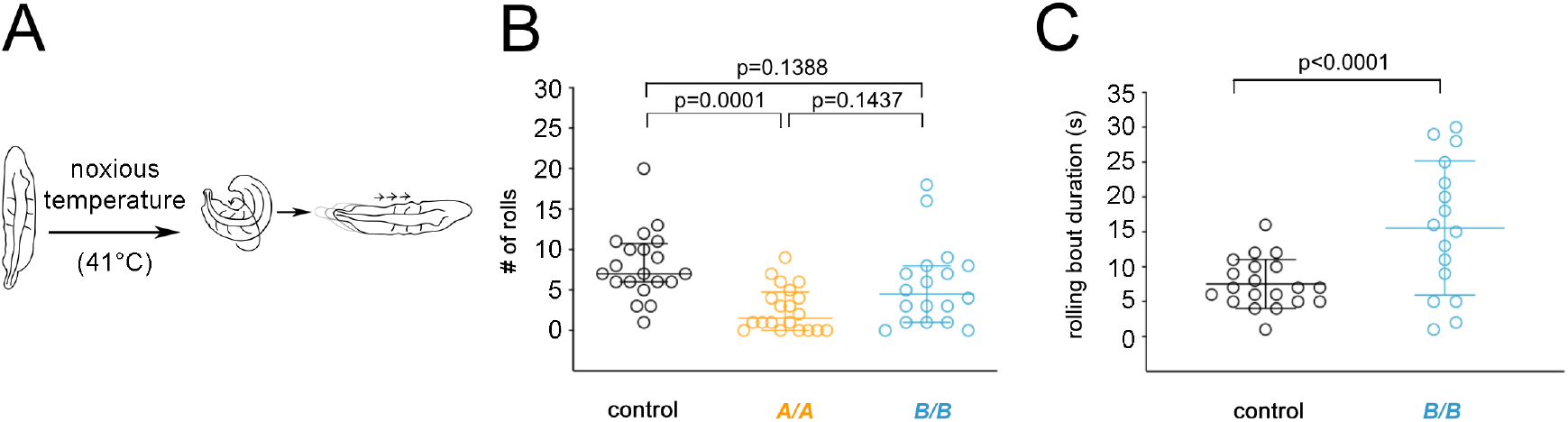
Alternative splicing of Dscam2 is required for robust behavioral responses to noxious stimuli. (A) Schematic of typical larval behavioral responses to noxious temperature. At 41 °C, larvae (left) initiate and bend and roll (middle) followed by rapid forward crawling (right). (B-C) Quantification of rolling behavior in control (*w^1118^), Dscam2A* single isoform homozygotes (*Dscam2^A/A^*) and *Dscam2B* single isoform homozygotes (*Dscam2^B/B^*). (B) *Dscam2^A/A^* mutants perform significantly fewer rolls than control. *w^1118^* vs. *Dscam2^A/A^*, p=0.0001; *w^1118^* vs. *Dscam2^B/B^*, p=0.1388; *Dscam2^A/A^* vs. *Dscam2^B/B^*, p=0.1437, as determined by Kruskal-Wallis test with Dunn’s multiple comparison test. Each dot is one larva, and median ± interquartile ranges are shown. (C) *Dscam2^B/B^* larvae do not perform significantly fewer rolls, but rolling bout duration was significantly increased compared to control larvae (p<0.0001 as determined by two-tailed unpaired t test). Each dot is one larva, and mean ± s.d. are shown.

## Discussion

In this study, we show that Dscam2A and 2B isoforms are expressed in a cell-type-specific pattern in the larval somatosensory circuit and that regulated alternative splicing is essential for nociceptive circuit wiring and behavior. Dscam2B is expressed in larval nociceptive sensory neurons, while Dscam2A is expressed in several central synaptic targets of nociceptive neurons. Artificially fixing matching isoforms in presynaptic and postsynaptic partners disrupted axon terminal morphology, including changes in connectives and reductions in presynaptic puncta.

Dscam2 isoforms bind homophilically, which is thought to trigger repulsion (Lah et al., 2014; Millard et al., 2007). In the visual system, cell-type specific Dscam2 isoform expression ensures that like-type axons are evenly spaced, whereas non-like-type axons can arborize in the same medulla column (Lah et al., 2014). Our data indicate that another role for regulated alternative splicing of a repulsive cell adhesion molecule such as Dscam2 is to ensure that pre- and postsynaptic partners do not express the same isoform. While splicing of surface receptors can have an instructive effect on axon guidance (Leggere et al., 2016) or selective trans-synaptic adhesion (Chih et al., 2006), we propose that regulated expression of Dscam2 isoforms provides a baseline repulsion between arbors that biases their connections. Regulated Dscam2 isoform expression guides their connections away from like-type developing neurites that express the same isoform and toward neurites with complementary isoform expression. Establishing such a baseline may be an early step in organizing axon topography where correct partners can readily interact. The various Dscam2 phenotypes in the nociceptive circuit and in other systems, tend to involve local changes in dendrite or axon patterning (Kerwin et al., 2018; Lah et al., 2014; Millard et al., 2007). These results are consistent with the idea that Dscam2 supports neural circuit connectivity at a local level, complementing instructive mechanisms that match synaptic partners.

Cell-type specific isoform expression of Dscam2B in sensory neurons can be controlled by the COE transcription factor Knot. Misexpression of Knot induced Dscam2B expression in one specific sensory neuron subtype, indicating that Knot can induce Dscam2B expression selectively in certain cellular contexts. Paradoxically, overexpression of Knot reduced Dscam2B expression in cIV neurons that normally express the Dscam2B isoform. Thus, Knot levels appear to be important for cell type-specific expression of Dscam2B. The RNA binding protein Muscleblind controls the mutually-exclusive choice between A and B isoforms by promoting Dscam2B and repressing Dscam2A expression (Li and Millard, 2019). Future studies may further examine the links between cell-type identity specification and specific Dscam2 isoform expression patterns in somatosensory, visual, and motor circuits.

Both Dscam2 single isoform homozygotes (*Dscam2^A/A^ and Dscam2^B/B^*) have defects in nocifensive rolling in addition to abnormal nociceptive sensory axon patterning, while *Dscam2^A/A^* and *Dscam2^B/B^* larvae showed different nocifensive rolling defects. Indeed, *Dscam2^A/A^* flies have stronger synaptic and morphological L1 and L2 phenotypes in the visual system than *Dscam2^B/B^* flies (Kerwin et al., 2018) and Dscam2B has an isoform-specific role in regulating synaptic strength in motor neuron boutons (Odierna et al., 2020). Dscam2 isoforms differ only by a single extracellular domain, which determines isoform recognition (Millard et al., 2007), however, how isoform-specific role is achieved for Dscam2 genes is unknown. Our results further support the notion that Dscam2 isoforms have distinct properties depending on the cell type in which they are expressed.

Our data indicate that regulated Dscam2 isoform expression is important for nociceptive circuit function, supporting the notion that regulated alternative splicing is critical for nervous system wiring and function. Our study adds to a series of Dscam2 studies that demonstrated how regulated alternative splicing of a gene contributes to the diverse aspects of the neurodevelopment, and that the morphological and synaptic wiring defects observed in Dscam2-deficient neurons have direct functional consequences (Bosch et al., 2015; Kerwin et al., 2018; Odierna et al., 2020). Future studies of Dscam2 in the nociceptive sensorimotor system should address the causal link between behavioral deficits and axon wiring and synapse phenotypes. Furthermore, due to the broad expression of Dscam2 in the nervous system and abundant tools for cell-type specific manipulation, this system may provide a useful model to further decipher the function of regulated alternative splicing in the nervous system.

## Materials and Methods

### Fly strains and maintenance

*Drosophila melanogaster* were reared using standard methods. Experimental crosses were maintained at 25°C. Third instar larvae were used. The following strains were used:

*w^1118^; UAS-CD8-GFP; Dscam2A-Gal4* (Lah et al., 2014)*; UAS-CD8-GFP; Dscam2B-Gal4* (Lah et al., 2014); *Dscam2A-LexA, LexAop-nucGFP* (Tadros et al., 2016); *Dscam2B-LexA, LexAop-nucGFP (Tadros et al., 2016); UAS-CD8-mCherry; 13XLexAop2-mCD8-GFP (BL32205); 412-Gal4 (Burgos et al., 2018); R72F11-Gal4* (BL39786); *ppk-CD4-tdTomato* (Han et al., 2011); *ppk-EGFP^1^* (Grueber et al., 2003); *Dscam2^A-A^/TM6B (Lah et al., 2014); Dscam2^B-B^/TM6B* (Lah et al., 2014); *UAS-Dscam2B (Millardet al., 2007); UAS-Dscam2A (Millardet al., 2007); hsFLP* (Gift from Gary Struhl); *13XlexAop2 (FRTstop) myr::smGdP-V5* (BL62107); *R38A10-LexA* (BL54106); *8XLexAop-Brpshort-cherry* (Berger-Muller et al., 2013); *13XLexAop2-IVS-myr-GFP (BL32212); ppk1.9-Gal4* (Ainsley et al., 2003); *ppk-CD4-tdGFP* (Han et al., 2011)

### Immunohistochemistry

Third instar larvae were dissected in 1 ×PBS, fixed in 4% paraformaldehyde (Electron Microscopy Sciences) in 1 × PBS for 15 minutes, rinsed for 15 minutes in 1 × PBS + 0.3% Triton X-100 (Sigma Aldrich) (PBS-TX) and blocked for at least 1 hour in 5% normal donkey serum (Jackson Immunoresearch) at 4°C. Primary antibodies used were chicken anti-GFP (1:1000; Abcam), rabbit anti-DsRed (1:200-1:500, Clontech), mouse anti-1D4 anti-Fasciclin II (1:10; Developmental Studies Hybridoma Bank), Goat anti-HRP (1:250, Jackson) and Rabbit anti-V5 Dylight 549 (1:250, Rockland). Larvae were incubated for 1-2 days in primary antibodies at 4°C, washed for at least 60 minutes in PBS-TX, and incubated 1-2 days in 4°C in species-specific, fluorophore-conjugated secondary antibodies (Jackson ImmunoResearch) at 1:200 in PBS-TX. Tissue was mounted on poly-L-lysine coated coverslips, dehydrated in ethanol series, cleared in xylenes, and mounted in DPX (Electron Microscopy Sciences). For FLPouts, embryos were collected for 1-3 days at 25°C, heat shocked at 38°C for 4-8 minutes, raised at 25°C until dissection at third instar stage.

### Behavior

Larvae were rinsed briefly with distilled water to remove food particles. Larvae were placed on a 1% agarose gel with 0.6% black ink (Higgins Waterproof Drawing Ink) heated to ~41°C (TE Technology Peltier; CP-031 and temperature controller; TC-36-24-RS232). Larvae were recorded on a Leica M50 camera with Leica FireCam software and QuickTime screen capture for 30 seconds. Videos were blinded and quantified offline. Larvae were tested one at a time and each animal was tested once. Larvae were scored for total number of rolls performed in a 30 second trial and the duration of the rolling bout, which was defined as the start of the first completed roll to the end of the last completed roll.

### Image Acquisition

Images were acquired on Zeiss 510 Meta and Zeiss 700 confocal microscopes with LSM software (Zeiss) using 40× Plan Neofluar objective. Image analysis was done using FIJI (Schindelin et al., 2012). Images with Brp-short labeling were processed using an ImageJ macro (Galindo et al., 2023).

### Quantification and Statistical Analysis

#### Statistics

Statistical details for each experiment can be found in figure legends. Data were tested for normality using the Shapiro-Wilk test. For normally distributed data, unpaired t-tests were used to compare two groups, and ordinary one-way ANOVA with Sidak’s multiple comparison tests were used to compare more than two groups. For non-normal data, Mann-Whitney U tests were used to compare two groups and Kruskal-Wallis with Dunn’s multiple comparisons test were used when comparing more than two groups. For categorical data, a two-tailed Fisher’s exact test was used. Uppercase (N) refers to individual larva and lowercase (n) refers to individual cells. Statistical tests were performed in GraphPad Prism. Exact p-values are displayed in Figures and listed in Figure Legends.

#### class IV axon scaffold missing longitudinals

For each larva, the total number of breaks in the axon scaffold were counted. Axons from segments A2-A7 were analyzed. A break was defined as discontinuous axons along the A-P axis.

#### class IV FLPout analysis

Each class IV axon terminal was scored for number of longitudinal branches. ‘2 branches’ indicate that the axon extended an anterior and posterior branch, ‘1 branch’ indicates that the axon extended either an anterior or posterior branch, and ‘0 branch’ indicates that no longitudinal branches were observed.

#### class IV axon defasciculation

Axons from segments A2-A7 were analyzed. For each neuromere, the total number of segregated longitudinal branches were counted. For example, axons that formed a cohesive bundle would be scored as 1 whereas axons that were segregated would be scored as >1, representing the number of visible cohesive bundles forming longitudinal branches for each neuromere.

#### Brp-short fluorescence

Axons from segments A4-A7 were analyzed. An ROI surrounding cIV axon terminals was generated by applying a threshold to the cIV channel and using the Analyze Particles command to create an outline. The ROI was overlaid to the Brp-short channel and the sum of pixels was measured.

## Acknowledgments

We thank Larry Zipursky for sharing fly stocks. We thank past and present members of the Grueber lab for discussion and feedback. We thank Zuckerman Institute’s Cellular Imaging platform for technical advice. Stocks in this study were obtained from the Bloomington Drosophila Stock Center. The monoclonal antibody 1D4 anti-Fasciclin II, developed by C. Goodman, was obtained from the Developmental Studies Hybridoma Bank, created by the NICHD of the NIH and maintained at The University of Iowa, Department of Biology, Iowa City, IA 52242.

## Competing interests

No competing interests declared.

## Funding

Research reported in this publication was supported by the National Institute of Neurological Disorders and Stroke of the National Institutes of Health under Award Number R01NS061908 (W.B.G.). This work was also supported by the National Institutes of Health NS098765 (S.E.G.). The content is solely the responsibility of the authors and does not necessarily represent the official views of the National Institutes of Health.

## Data Availability

All relevant data can be found within the article.

## Author contributions

Conceptualization: W.B.G, S.E.G, and G.J.S.; Methodology: S.E.G., G.J.S.; Formal analysis: S.E.G., G.J.S.; Investigation: S.E.G.; Writing – original draft: W.B.G., S.E.G., G.J.S.; Writing – reviewing & editing: W.B.G., S.E.G., G.J.S., S.S.M.; Visualization: S.E.G.; Resources: S.S.M.; Supervision: W.B.G.; Funding acquisition: W.B.G., S.E.G.

## Notes

### Competing Interest Statement

The authors have declared no competing interest.

